## Supplemental Methods and Results for "Coevolution of relative brain size and life expectancy in parrots"

31-12-2021

**Supplemental Results** **9**

**References** **13**

### Supplemental Methods

#### Standardisation taxonomy

All species names were converted to species-level names using the taxonomy of the IUCN Red List of Threatened Species$^{TM}$ (IUCN 2020) prior to any analysis. If the species names were not found as accepted species names or in the synonym list, fuzzy matching was attempted using the function *stringdist* from the package *stringdist* (van der Loo 2014) in R (R Core Team 2021) with default settings. A match was accepted if the distance was less than four characters. Species that were still not found were manually matched (if possible) using the IUCN website and Avibase (Lepage, Vaidya, and Guralnick 2014). If records were at subspecies level, taxonomy was standardised using a subspecies level list from IUCN (IUCN 2020), which assigns subspecies to their respective parent taxon. If the name was not found there the species level name was used for standardisation. A full translation list can be found in the Supplemental Materials.

#### Life expectancy pre-processing

We extracted data on birth and death dates from the Species360 Zoological Information Management System (ZIMS) (Species360 2020). We only included a total number of 244 species with records for at least 50 individuals to ensure sufficient data for a reliable fit of life expectancy. We ran BaSTA models on a random subset for species with large sample size (N > 1000) to determine this threshold. We only included individuals that died after 1980, when husbandry guidelines were greatly improved. We further excluded individuals with lifespans that were outside the 99th percentile of the species lifespan to minimise the number of data entry errors. We also excluded stillbirths (i.e., individuals that died on the day of birth), individuals of unknown sex and surgically contracepted individuals. For the estimation of life expectancies see the main text.

#### Body weight

We extracted weight measurement from ZIMS and only included captive born individuals where the birth date was not estimated. We also only included individuals for which the lifespan was within the 99.9% range to minimize the potential data-entry errors. Only measurements with a 'normal' health status were included. Clear outliers with respect to age and weight were removed manually to reduce the effect of data-entry errors. Only weight measurements of individuals older than half the maximum age (measured on a log scale) were included. This threshold was chosen such that only adult weights were included. Since there is no consensus on physical maturity for all species and fitting a bespoke growth curve was outside the scope of this paper, this was the best available method to exclude juveniles. Additional outliers were removed if they were larger than the third quantile plus three times the inter-quantile distance or smaller than the first quantile minus three times the inter-quantile distance (on a log scale). All measurements were plotted per species and the cleaning performance was assessed visually.

Additional measurements were included from the literature for species where no ZIMS data were available (Conde et al. 2019). For these species the value from the literature was included as a single measurement for a single individual since no sample size or standard error was reported in most cases.

To estimate the average adult body weight per species we ran a Bayesian model with the following structure:

$$
\begin{aligned}
\text{log weight}_i &\sim \text{normal}(\mu_i, \sigma) \\
\mu_i &= \alpha_{\text{species}[i]} + z_{\text{individual}[j]} * \phi_{\text{species}[i]} \\
\alpha_{\text{species}} &\sim \text{normal}(6, 6) \\
z_{\text{individual}} &\sim \text{normal}(0, 1) \\
\phi_{\text{species}} &\sim \text{lognormal}(\overline{\phi}, s) \\
\overline{\phi} &\sim \text{normal}(-2, 0.5) \\
s &\sim \text{exponential}(5) \\
\sigma &\sim \text{exponential}(1)
\end{aligned}
$$

Where $\mu_i$ is the species-level average, $\alpha_{\text{species}[i]}$ the species-level intercept, $z_{\text{individual}[j]}$ the z-score of the individual-level off-set, $\phi_{\text{species}[i]}$ the species-level standard deviation of the off-set, $\overline{\phi}$ the average standard deviation of the individual-level off-set, $s$ the standard deviation of the aforementioned and $\sigma$ the standard deviation of the distribution of log weights. Priors were chosen with prior predictive simulations to ensure that predictions were mostly restricted to reasonable values. The multi-level prior for the individual variation was used to inform parameters for species with only a single individual. The model was run using the function *ulam* from the package *rethinking* (McElreath 2020) in R, which is an interface to run the No U-turn Sampler (an improved version of the Hamiltonian Monte Carlo algorithm) in Stan (Gelman, Lee, and Guo 2015). The estimated means and standard errors of the mean were exported and used in further analysis.

#### Brain size

Raw brain size measurements were compiled from three sources: new measurements of brain weight, brain weight taken from Iwaniuk, Dean, and Nelson (2005) and endocranial volumes taken from Ksepka et al. (2020). Brain weight was estimated from endocranial volumes using: 1.036 g/ml (Kretschmann and Wingert 1969). Iwaniuk and Nelson (2002) showed that endocranial volumes are highly correlated with whole brain weight in birds and that bias should only be a problem when looking at brain areas or seasonal variation. To estimate the average log adult brain weight per species we ran a Bayesian model with the following structure:

$$
\begin{aligned}
\text{brain weight}_i &\sim \text{normal}(\mu_i,\ \sigma) \\
\mu_i &= \alpha_{\text{species}[i]} \\
\alpha_{\text{species}} &\sim \text{normal}(\overline{\alpha},\ \text{s}) \\
\overline{\alpha} &\sim \text{normal}(2,\ 1) \\
\text{s} &\sim \text{exponential}(1) \\
\sigma &\sim \text{exponential}(1)
\end{aligned}
$$

Priors were chosen with prior predictive simulations to ensure that predictions were mostly restricted to reasonable values. We choose to run a multi-level model to inform the overall mean ($\overline{\alpha}$) by the data rather than setting it to a fixed value. The model was run using the function *ulam* from the package *rethinking* in R. The estimated means and standard errors of the mean were exported and used in further analysis. After estimation of averages from raw data, we added additional summarised data from Schuck-Paim, Alonso, and Ottoni (2008) estimating SE from the reported sample sizes and the standard variation from the raw data.

#### Diet

We compiled diet data from the primary literature and by contacting individual researchers. The reports from experts working specifically on a particular species were given priority. We scored diet composition across nine categories, which were either considered to have high or low protein content (low: nectar, sugar/honeydew, fruit, leaves/shoots/buds trees, leaves/shoots/buds ground cover, bulbs/roots; high: seeds, nuts, meat). For each category, a species could use it as either a main food source (multiple possible), an extra food source or rarely/never consume it. The final score was then: 1) high protein, if the species only had high protein sources as main diet; 2) medium protein, if the species had main sources across high and low protein categories; or 3) low protein, if the species only had low protein sources as main diet. We treated diet as an ordinal variable.

#### Insularity

We considered species insular if their native range only included non-continental ranges. We compiled this variable from Sayol et al. (2019), Ducatez et al. (2020) and IUCN (2020). We treated this variable as binomial.

#### Latitude

To account for seasonal variability we included the most extreme range (furthest from the equator) that a species inhabits. We downloaded all ranges included in Billerman et al. (2020), which is the same source used for IUCN threat-status assessments. We only included native ranges and took the maximum absolute latitude (e.g. furthest distance from the equator) for each species.

#### Developmental time

Developmental time was included as incubation period + fledging period. Both these variables were compiled from Amniote Life History Database (Myhrvold et al. 2015) (ALHD) and several handbooks and papers Del Hoyo et al. (1997). ALHD data was preferred over other sources. If multiple handbooks had values, the mean was used. We log transformed the data to get an approximately normally distributed variable.

#### Clutch size

Clutch size was compiled from the ALHD, Demographic Species Knowledge Index (Conde et al. 2019) (D-SKI) and several handbooks and papers Del Hoyo et al. (1997). Because data from D-SKI already included data from the ALHD, D-SKI data was preferred over ALHD over other sources. If data was available from multiple handbooks we used the mean value. We log transformed the data to get an approximately normally distributed variable.

#### Age of first reproduction

For age of first reproduction (AFR) we used parentage records from ZIMS. Since parrots were sometimes kept without a bird of the opposite sex, we calculated at what age 5% of the population had produced a live offspring for the first time. We only did this for species that had at least 30 individuals that had reproduced. We only included individuals that were captive born and had a known birth date. We removed cases where AFR was below 6 months or above 80 years, as these were clearly data entry errors. We calculated this proxy of AFR for 90 species and the log standardised values in the third model.

#### Full statistical models

We ran three models to test which hypothesis best explained the observed data. All models were implemented using *ulam* from the *rethinking* package. We specified a structural equations model with a main model that explain life expectancy given a set of covariates. Each of the covariates had a sub-model that explained this variable given the relationships from the DAG (see main Figure 1). We also included a phylogenetic variance-covariance for all variables and standard error for life expectancy, body mass and brain mass.

##### Model 1

Model 1 only included body mass, relative brain mass, latitude and diet as covariates The structure was as follows:

main model:

$$\mathrm{LE}_{\mathrm{obs},i} \sim \mathrm{normal}(\mathrm{LE}_{\mathrm{true},i},\ \mathrm{SE}_{\mathrm{LE},i})$$ LE = standardised life expectancy

$$\mathbf{LE}_{\mathrm{true}} \sim \mathrm{MVnormal}(\boldsymbol{\mu}_{\mathrm{LE}},\ \mathbf{S}_{\mathrm{LE}})$$ SE = standard error

$\boldsymbol{\mu}_{\mathrm{LE}}$ = mean life expectancy per species

*obs* = observed data

*true* = modelled underlying true values

$\mathbf{S}$ = variance co-variance matrix

$$\mu_{\mathrm{LE},i} = \alpha_{\mathrm{I}[i]} + \beta_{\mathrm{BO}} * \mathrm{BO}_{\mathrm{true},i} + \beta_{\mathrm{BR}} * (\mathrm{BR}_{\mathrm{true},i} - \mu_{\mathrm{BR},i}) + \beta_{\mathrm{LA}} * \mathrm{LA}_i + \beta_{\mathrm{PD}} * \sum_{j=0}^{\mathrm{PD}_i - 1} \delta_{\mathrm{PD},j}$$

I = index variable for insularity

BO = standardised body mass

BR = standardised brain mass

LA = standardised maximum latitude

D = protein level diet

$\delta$ = marginal diet effect

priors main model:

$$\boldsymbol{\alpha},\ \beta_{\mathrm{BO}},\ \beta_{\mathrm{BR}},\ \beta_{\mathrm{I}},\ \beta_{\mathrm{LA}},\ \beta_{\mathrm{PD}} \sim \mathrm{normal}(0,\ 0.5)$$

$$\boldsymbol{\delta}_{\mathrm{PD}} \sim \mathrm{dirichlet}(2,\ 2)$$

$$\mathrm{S}_{\mathrm{LE},ij} = \eta^2_{\mathrm{LE}} \exp(-\rho^2_{\mathrm{LE}}\ \mathrm{P}^2_{ij}) + \delta_{\mathrm{P},ij}\ \sigma^2_{\mathrm{P}}$$ formula for L2 norm

$$\eta^2_{\mathrm{LE}} \sim \mathrm{exponential}(2)$$ P = normalised phylogenetic distance

$$\rho^2_{\mathrm{LE}} \sim \mathrm{exponential}(0.1)$$

Sub-models in model 1:

sub-model latitude:

$$\begin{aligned}
\mathbf{LA} &\sim \text{MVnormal}(\boldsymbol{\mu}_{\text{LA}},\ \mathbf{S}_{\text{LA}}) \\
\mu_{\text{LA},i} &\sim \text{normal}(0,\ 0.5) \\
\text{S}_{\text{LA},ij} &= \eta^2_{\text{LA}}\ \exp(-\rho^2_{\text{LA}}\ \text{P}^2_{ij})\ +\ \delta_{\text{P},ij}\ \sigma^2_{\text{P}} \\
\eta^2_{\text{LA}} &\sim \text{exponential}(2) \\
\rho^2_{\text{LA}} &\sim \text{exponential}(0.1)
\end{aligned}$$

sub-model relative brain size:

$$\begin{aligned}
\text{BR}_{\text{obs},i} &\sim \text{normal}(\text{BR}_{\text{true},i},\ \text{SE}_{\text{BR},i}) \\
\mathbf{BR}_{\text{true}} &\sim \text{MVnormal}(\boldsymbol{\mu}_{\text{BR}},\ \mathbf{S}_{\text{BR}}) \\
\mu_{\text{BR},i} &= \xi\ +\ \phi * \text{BO}_{\text{true},i} \\
\xi,\ \phi &\sim \text{normal}(0,\ 0.5) \\
\text{S}_{\text{BR},ij} &= \eta^2_{\text{BR}}\ \exp(-\rho^2_{\text{BR}}\ \text{P}^2_{ij})\ +\ \delta_{\text{P},ij}\ \sigma^2_{\text{P}} \\
\eta^2_{\text{BR}} &\sim \text{exponential}(2) \\
\rho^2_{\text{BR}} &\sim \text{exponential}(0.1)
\end{aligned}$$

sub-model body mass:

$$\begin{aligned}
\text{BO}_{\text{obs},i} &\sim \text{normal}(\text{BO}_{\text{true},i},\ \text{SE}_{\text{BO},i}) \\
\mathbf{BO}_{\text{true}} &\sim \text{MVnormal}(\boldsymbol{\mu}_{\text{BO}},\ \mathbf{S}_{\text{BO}}) \\
\text{S}_{\text{BO},ij} &= \eta^2_{\text{BO}}\ \exp(-\rho^2_{\text{BO}}\ \text{P}^2_{ij})\ +\ \delta_{\text{P},ij}\ \sigma^2_{\text{P}} \\
\eta^2_{\text{BO}} &\sim \text{exponential}(2) \\
\rho^2_{\text{BO}} &\sim \text{exponential}(0.1)
\end{aligned}$$

To minimise confusion, conventional Greek symbols were used in combination with subscripts to denote to which part of the model they belong. Relative brain size was computed using the average expected brain mass ($\mu_{\text{BR},i}$) from a sub-model that run simultaneously. We used $\xi$ and $\phi$ for the intercept and slope of brain mass sub-model to avoid confusion the body mass effect on life expectancy with the effect on brain size. Missing values for life expectancy, brain mass, body mass and latitude were imputed using the above distributions.

**Model 2**

Model 2 also included developmental time and clutch size as predictors, but the structure of the other parts was the same. The main model was as follows:

main model:

$$\begin{aligned}
\mathrm{LE}_{\mathrm{obs},i} &\sim \mathrm{normal}(\mathrm{LE}_{\mathrm{true},i},\ \mathrm{SE}_{\mathrm{LE},i}) && \mathrm{LE} = \text{standardised life expectancy}\\
\mathbf{LE}_{\mathrm{true}} &\sim \mathrm{MVnormal}(\boldsymbol{\mu}_{\mathrm{LE}},\ \mathbf{S}_{\mathrm{LE}}) && \mathrm{SE} = \text{standard error}\\
& && \boldsymbol{\mu}_{\mathrm{LE}} = \text{mean life expectancy per species}\\
& && obs = \text{observed data}\\
& && true = \text{modelled underlying true values}\\
& && \mathbf{S} = \text{variance co-variance matrix}\\
\mu_{\mathrm{LE},i} &= \alpha_{\mathrm{I}[i]} + && \mathrm{I} = \text{index variable for insularity}\\
&\quad \beta_{\mathrm{BO}} * \mathrm{BO}_{\mathrm{true},i} + && \mathrm{BO} = \text{standardised body mass}\\
&\quad \beta_{\mathrm{BR}} * (\mathrm{BR}_{\mathrm{true},i} - \mu_{\mathrm{BR},i}) + && \mathrm{BR} = \text{standardised brain mass}\\
&\quad \beta_{\mathrm{LA}} * \mathrm{LA}_i + && \mathrm{LA} = \text{standardised maximum latitude}\\
&\quad \beta_{\mathrm{PD}} * \sum_{j=0}^{\mathrm{PD}_i - 1} \delta_{\mathrm{PD},j} + && \mathrm{PD} = \text{protein level diet}\\
&\quad \beta_{\mathrm{DT}} * \mathrm{DT}_{\mathrm{true},i} + && \delta = \text{marginal diet effect}\\
&\quad \beta_{\mathrm{CS}} * \mathrm{CS}_{\mathrm{true},i} + && \mathrm{DT} = \text{developmental time}\\
& && \mathrm{CS} = \text{clutch size}
\end{aligned}$$

priors main model:

$$\begin{aligned}
\boldsymbol{\alpha},\ \beta_{\mathrm{BO}},\ \beta_{\mathrm{BR}},\ \beta_{\mathrm{I}},\ \beta_{\mathrm{LA}},\ \beta_{\mathrm{PD}},\ \beta_{\mathrm{DT}},\ \beta_{\mathrm{CS}} &\sim \mathrm{normal}(0,\ 0.5)\\
\boldsymbol{\delta}_{\mathrm{PD}} &\sim \mathrm{dirichlet}(2,\ 2)\\
\mathrm{S}_{\mathrm{LE},ij} &= \eta^2_{\mathrm{LE}} \exp(-\rho^2_{\mathrm{LE}}\ \mathrm{P}^2_{ij}) + \delta_{\mathrm{P},ij}\ \sigma^2_{\mathrm{P}} && \text{formula for L2 norm}\\
\eta^2_{\mathrm{LE}} &\sim \mathrm{exponential}(2) && \mathrm{P} = \text{normalised phylogenetic distance}\\
\rho^2_{\mathrm{LE}} &\sim \mathrm{exponential}(0.1)
\end{aligned}$$

The additional sub-models in model 2:

$$\begin{aligned}
&\underline{\text{sub-model developmental time:}} \\
\mathbf{DT} &\sim \text{MVnormal}(\boldsymbol{\mu}_{\text{DT}},\ \mathbf{S}_{\text{DT}}) \\
\mu_{\text{DT},i} &= \zeta\ +\ \theta * \text{BO}_{\text{true},i}\ +\ \iota * (\text{BR}_{\text{true},i}\ - \mu_{\text{BR},i}) \\
\zeta,\ \theta,\ \iota &\sim \text{normal}(0,\ 0.5) \\
\text{S}_{\text{DT},ij} &= \eta^2_{\text{DT}}\ \exp(-\rho^2_{\text{DT}}\ \text{P}^2_{ij})\ +\ \delta_{\text{P},ij}\ \sigma^2_{\text{P}} \\
\eta^2_{\text{DT}} &\sim \text{exponential}(2) \\
\rho^2_{\text{DT}} &\sim \text{exponential}(0.1) \\
&\underline{\text{sub-model developmental time:}} \\
\mathbf{CS} &\sim \text{MVnormal}(\boldsymbol{\mu}_{\text{CS}},\ \mathbf{S}_{\text{CS}}) \\
\mu_{\text{CS},i} &= \kappa\ +\ \omega * \text{LA}_i\ +\ \lambda * (\text{BR}_{\text{true},i}\ - \mu_{\text{BR},i}) \\
\kappa,\ \omega,\ \lambda &\sim \text{normal}(0,\ 0.5) \\
\text{S}_{\text{CS},ij} &= \eta^2_{\text{CS}}\ \exp(-\rho^2_{\text{CS}}\ \text{P}^2_{ij})\ +\ \delta_{\text{P},ij}\ \sigma^2_{\text{P}} \\
\eta^2_{\text{CS}} &\sim \text{exponential}(2) \\
\rho^2_{\text{CS}} &\sim \text{exponential}(0.1)
\end{aligned}$$

##### Model 3

To test if including AFR made a difference a third model was run. We sub-setted the data to species where we had AFR available and did not try to impute this variable. The structure of the sub models was the same as model 2, and we added $\beta_{\text{AFR}} * \text{AFR}_i$ as a predictor in the main model, where $\beta_{\text{AFR}} \sim \text{normal}(0,\ 0.5)$.

##### Priors

Priors for the intercept ($\alpha$) and slopes ($\beta$s) were chosen such that most of the prior assigned effects were between -1 and 1 with no effect as the most likely effect, thereby mildly regularising the model to conservative values. The prior for the delta variables was chosen to be mildly regularising as well. For the $\eta^2$ and $\rho^2$ priors we used prior predictive simulations to make sure that the covariance structure allowed by the prior stayed within possible bounds while still allowing for both very strong and no covariance.

### Supplemental Results

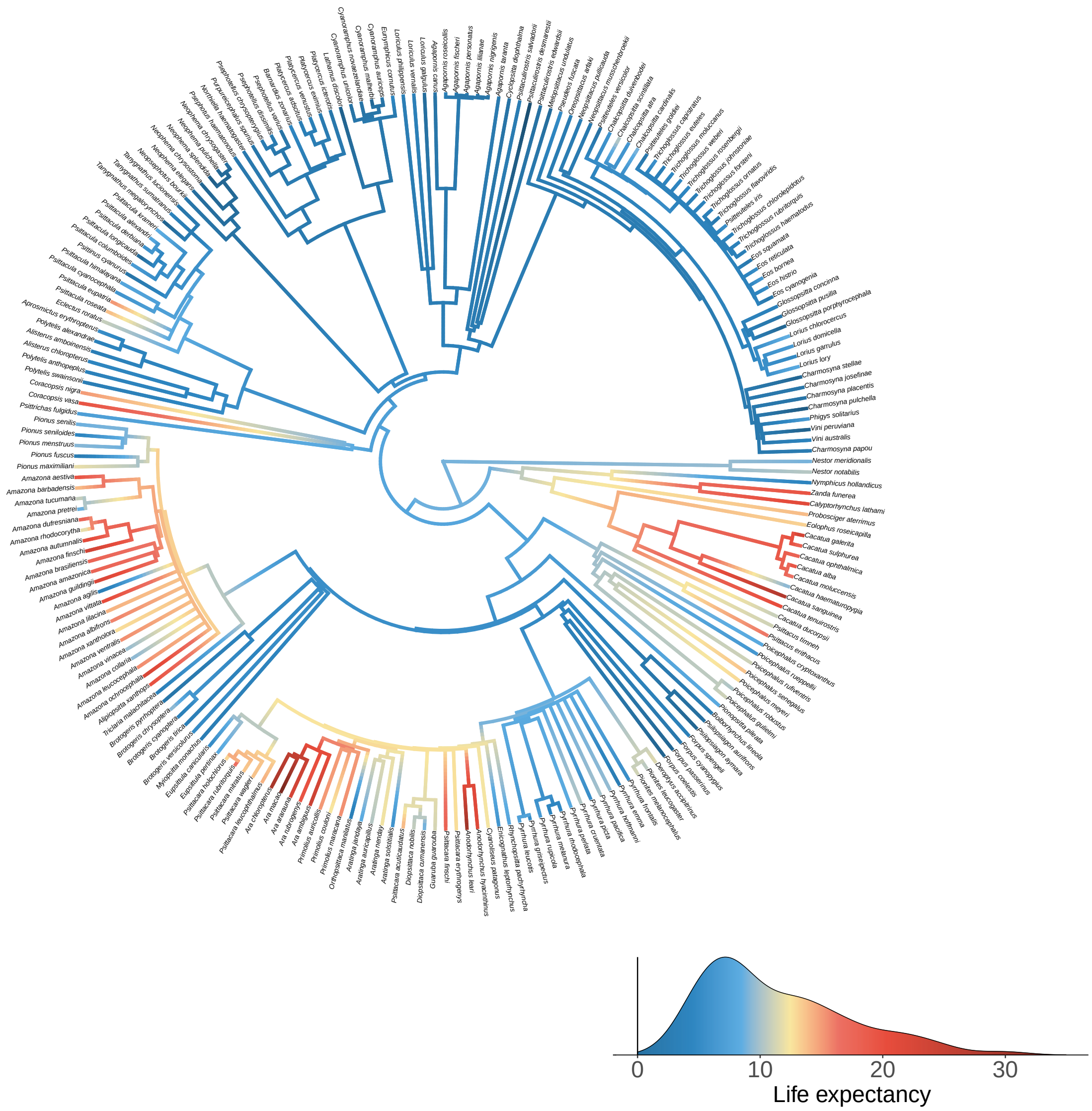

Figure S1: Phylogenetic tree of all species for which enough data was available. Branches are coloured acording to life expectancy (see density plot in bottom right).

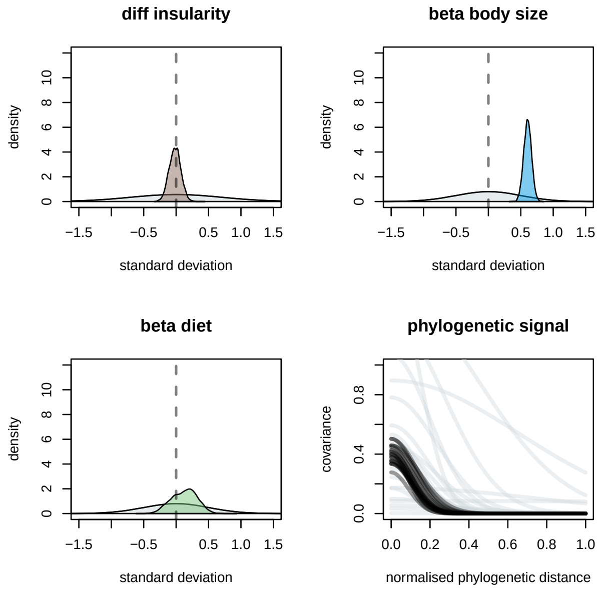

Figure S2: Slope estimates for predictor variables and phylogenetic signal in model 1. For insularity the difference between islandic and continental species is shown. Grey density plots and lines are the regularising priors. Coloured areas are the posterior densities for the parameters. Black lines are 20 samples of the posterior for the phylogenetic covariance.

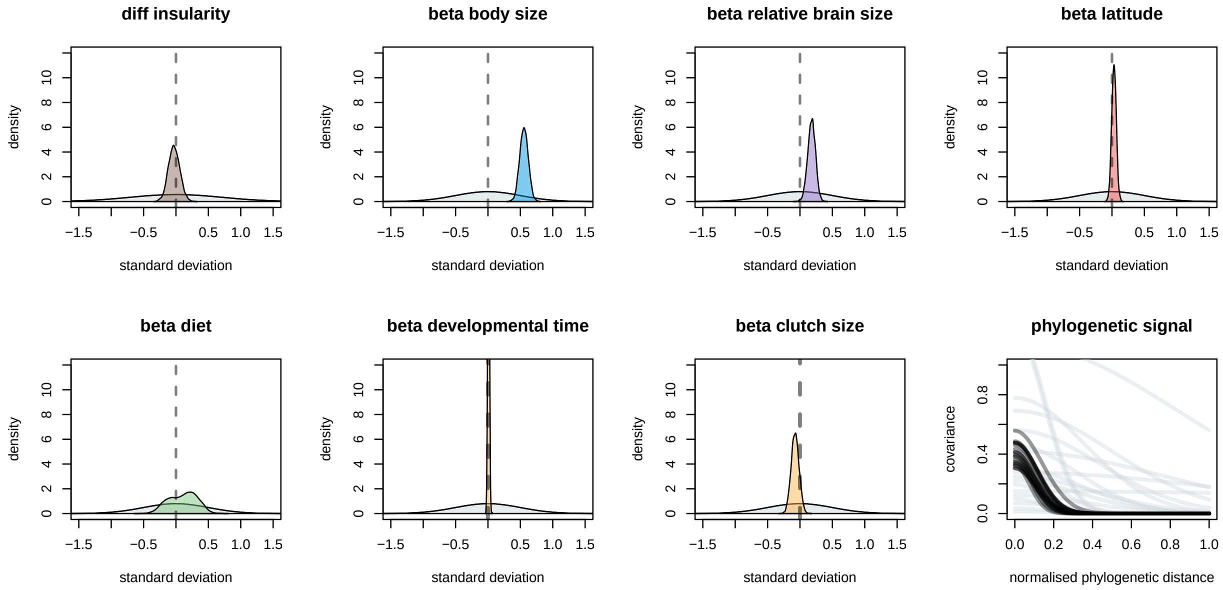

Figure S3: Slope estimates for predictor variables and phylogenetic signal in model 2. For insularity the difference between islandic and continental species is shown. Grey density plots and lines are the regularising priors. Coloured areas are the postior densities for the parameters. Black lines are 20 samples of the posterior for the phylogenetic covariance.

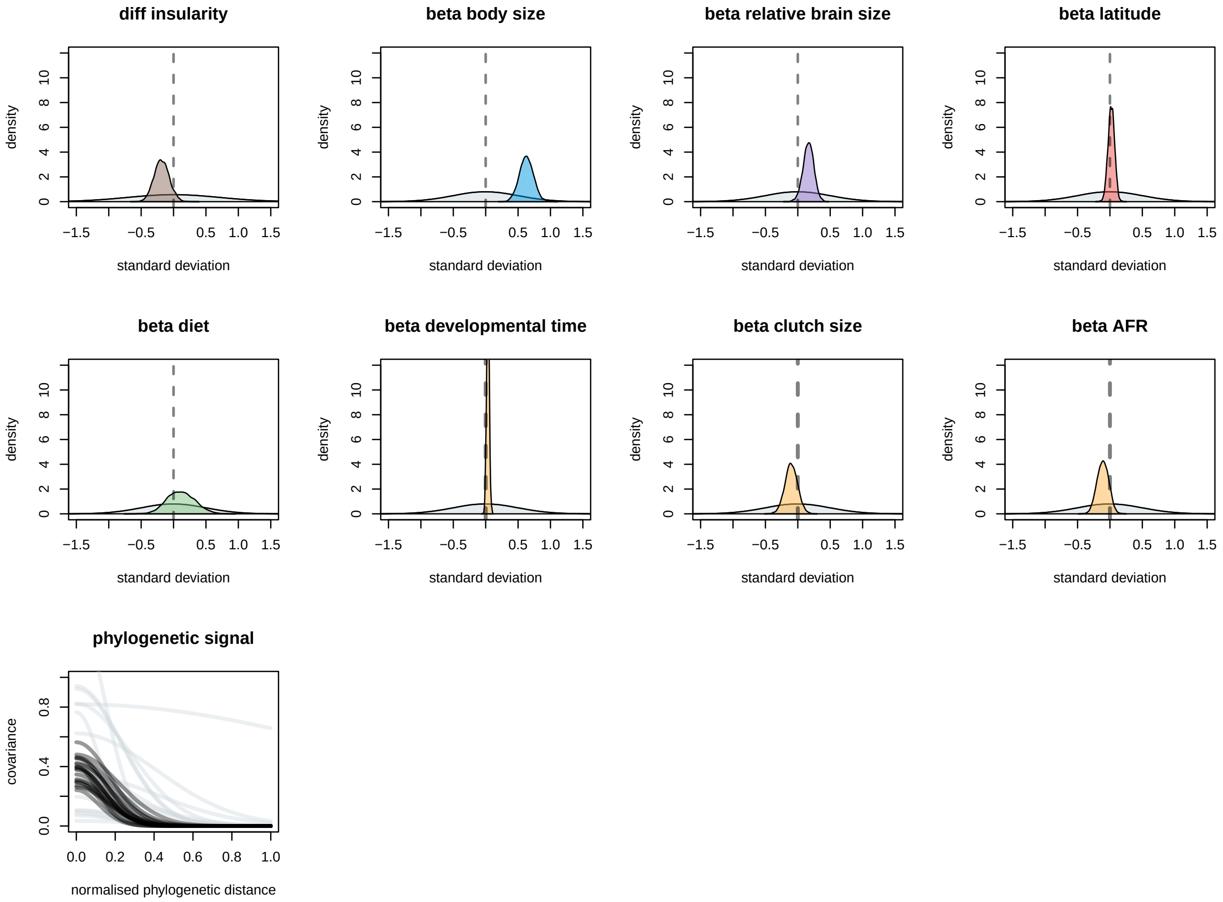

Figure S4: Slope estimates for predictor variables and phylogenetic signal in model 3. For insularity the difference between islandic and continental species is shown. Grey density plots and lines are the regularising priors. Coloured areas are the postior densities for the parameters. Black lines are 20 samples of the posterior for the phylogenetic covariance.

Forshaw, Joseph M. 2002. *Australian Parrots*. Alexander Editions.

———. 2010. *Parrots of the World*. Princeton University Press.

Gelman, Andrew, Daniel Lee, and Jiqiang Guo. 2015. "Stan: A Probabilistic Programming Language for Bayesian Inference and Optimization." *Journal of Educational and Behavioral Statistics* 40 (5): 530–43.

IUCN. 2020. "The IUCN Red List of Threatened Species." 2020. https://www.iucnredlist.org.

Iwaniuk, Andrew N, Karen M Dean, and John E Nelson. 2005. "Interspecific Allometry of the Brain and Brain Regions in Parrots (Psittaciformes): Comparisons with Other Birds and Primates." *Brain, Behavior and Evolution* 65 (1): 40–59.

Iwaniuk, Andrew N, and John E Nelson. 2002. "Can Endocranial Volume Be Used as an Estimate of Brain Size in Birds?" *Canadian Journal of Zoology* 80 (1): 16–23.

Kretschmann, H-J, and F Wingert. 1969. "Biometrische Analyse Der Volumina Des Striatum Einer Ontogenetischen Reihe von Albinomäusen." *Zeitschrift für Anatomie Und Entwicklungsgeschichte* 128 (2): 85–108.

Ksepka, Daniel T, Amy M Balanoff, N Adam Smith, Gabriel S Bever, Bhart-Anjan S Bhullar, Estelle Bourdon, Edward L Braun, et al. 2020. "Tempo and Pattern of Avian Brain Size Evolution." *Current Biology* 30 (11): 2026–36.

Lepage, Denis, Gaurav Vaidya, and Robert Guralnick. 2014. "Avibase–a Database System for Managing and Organizing Taxonomic Concepts." *ZooKeys*, no. 420: 117.

McElreath, Richard. 2020. *Rethinking: Statistical Rethinking Book Package*.

Myhrvold, Nathan P, Elita Baldridge, Benjamin Chan, Dhileep Sivam, Daniel L Freeman, and SK Morgan Ernest. 2015. "An Amniote Life-History Database to Perform Comparative Analyses with Birds, Mammals, and Reptiles: Ecological Archives E096-269." *Ecology* 96 (11): 3109–9.

R Core Team. 2021. *R: A Language and Environment for Statistical Computing*. Vienna, Austria: R Foundation for Statistical Computing. https://www.R-project.org.

Sayol, Ferran, Oriol Lapiedra, Simon Ducatez, and Daniel Sol. 2019. "Larger Brains Spur Species Diversification in Birds." *Evolution* 73 (10): 2085–93.

Schuck-Paim, Cynthia, Wladimir J Alonso, and Eduardo B Ottoni. 2008. "Cognition in an Ever-Changing World: Climatic Variability Is Associated with Brain Size in Neotropical Parrots." *Brain, Behavior and Evolution* 71 (3): 200–215.

Smiet, Fred. 1985. "Notes on the Field Status and Trade of Moluccan Parrots." *Biological Conservation* 34 (2): 181–94.

Species360. 2020. *Zoological Information Management System (ZIMS)*. https://www.zims.Species360.org.

van der Loo, M. P. J. 2014. “The Stringdist Package for Approximate String Matching.” *The R Journal* 6: 111–22. https://CRAN.R-project.org/package=stringdist.
